# SLCs contribute to endocrine resistance in breast cancer: role of SLC7A5 (LAT1)

**DOI:** 10.1101/555342

**Authors:** Catherine M. Sevigny, Surojeet Sengupta, Zhexun Luo, Xiaoyi Liu, Rong Hu, Zhen Zhang, Lu Jin, Dominic Pearce, Diane Demas, Ayesha N. Shajahan-Haq, Robert Clarke

## Abstract

Resistance to endocrine therapies remains a major challenge for the successful management of patients with estrogen receptor-positive (ER+) breast cancers. Central to the development of resistance is the adaptive reprogramming of cellular metabolism in response to treatment. Solute carriers (SLCs) play a key role in metabolic reprogramming by transporting sugars, amino acids, and other nutrients and regulating their abundance within the cell and its subcellular organelles. We found 109 SLC mRNAs to be differentially expressed between endocrine sensitive and resistant breast cancer cells. In univariate analyses, 55 of these SLCs were associated with poor outcome in ER+ breast cancer patients. Data from TMT and SILAC studies then led us to focus on SLC7A5 (LAT1). In complex with SLC3A2 (CD98), LAT1 is the primary transporter of large, neutral amino acids including leucine and tyrosine. LAT1 expression is estrogen-regulated in endocrine sensitive cells but this regulation is lost in resistant cells. Pharmacologic inhibition or genetic depletion of LAT1 each suppressed growth in two models of endocrine resistant breast cancer. Autophagy was activated with LAT1 inhibition, but cells failed to degrade p62 showing that flux was blocked. Overexpression of the LAT1 cDNA increased protein synthesis and high LAT1 expression correlated with poor disease-free survival in ER+ breast cancer patients. This study uncovers a novel LAT1 mediated adaptive response that contributes to the development of endocrine resistance. Blocking LAT1 function may offer a new avenue for effective therapeutic intervention against endocrine resistant ER+ breast cancers.

## Introduction

In the United States, breast cancer is the most commonly diagnosed cancer in women^1^. Of the 253,000 newly diagnosed breast cancers each year, approximately 70% are estrogen receptor positive (ER+)^2^. Endocrine therapies, such as aromatase inhibitors (AIs) and selective estrogen receptor modulators (SERMs), have extended life expectancy for patients with ER+ disease^3^. Unfortunately, resistance to these treatments is common^4,5^. Patients who do not initially respond to endocrine therapies (*de novo* resistance), or who initially respond but eventually recur (acquired resistance), generally require cytotoxic chemotherapies. Chemotherapy often induces serious side effects^6^ but is rarely curative in advanced disease. It is critical to understand how resistance to endocrine therapy develops and to design more effective treatments for patients. Ideally, this can be achieved while minimizing toxicity.

Dysregulation of cellular energetics, a key hallmark of cancer, is driven by altered metabolism in cancer cells compared with normal cells^7,8^. Unique aspects of cancer cell metabolism can use pro-survival mechanisms, such as autophagy, to survive under stress or in a nutrient-poor microenvironment. Autophagy is an intracellular process of lysosomal degradation of proteins and organelles that can release amino acids, sugars, and other essential nutrients to support cell metabolism^9,10^ and help to meet cellular energy demand^11^. If proliferation does not resume and autophagy remains active at a high level, autophagy can switch from being pro-survival to activating cell death. Previously, we have shown that endocrine resistant cells exhibit a higher autophagy efficiency than sensitive cells^12^. Differential expression of proteins involved in metabolism likely contributes to maintaining the balance between pro-survival autophagy and pro-apoptotic responses to endocrine therapies^13^.

We used established models of endocrine resistant breast cancer to assess changes in the patterns of protein expression of sensitive (LCC1^14^; estrogen independent, tamoxifen sensitive) and resistant (LCC9^15^; estrogen independent, tamoxifen and fulvestrant cross resistant) cells^15^. We also used the T47D variants T47D:A18 (estrogen dependent, tamoxifen sensitive), T47D:A18-4HT^16^ (estrogen independent, tamoxifen resistant) and T47D:C42^17^ (estrogen receptor negative, tamoxifen resistant). Together, these models reflect the endocrine therapy sensitive and resistant phenotypes that exist in some patient cohorts. Differential mRNA expression analysis of endocrine sensitive (LCC1) and endocrine resistant (LCC9) cells implicated several solute carriers (SLCs) in acquired endocrine resistance. At the mRNA level, we found altered expression of 109 members of the SLC gene family; 16 of these genes were confirmed to be differentially expressed by unbiased proteome analyses. Two quantitative proteomic approaches were used to study differential protein expression of LCC1 and LCC9 cells: 1) tandem mass tag (TMT) and 2) stable isotope labeling with amino acids in cell culture (SILAC). We hypothesized that changes in solute carriers (SLCs) expression and nutrient uptake may supplement autophagy to support the cellular metabolism that drives an endocrine resistant phenotype.

SLCs are transport proteins that can act as exchangers, cotransporters, facilitated transporters, or orphan transporters for key nutrients such as amino acids or sugars^18^. Here we show that the solute carrier family 7 (SLC7) has several members upregulated in resistant compared with sensitive cells. SLC7s transport amino acids into cells and can feed intermediate metabolism^12,19^. Amino acids can be modified to enter the citric acid cycle, such as with the conversion of leucine into acetoacetate^20,21^. SLC7A5 (also known as LAT1) and/or its interacting partner SLC3A2 (CD98) is upregulated in a variety of cancers^22–26^ and is critical for growth and survival. Amino acids including the essential amino acid leucine and the non-essential amino acid tyrosine are transported into the cell through LAT1^23,27^. For example, upregulation of LAT1 during androgen therapy can drive pancreatic cancer progression^22^, whereas homozygous knockout of LAT1 is embryonic lethal in mice^28^. We chose to focus here on LAT1 because it was significantly upregulated in the LCC9 compared to the LCC1 cells in both proteome analyses and in the transcriptome analysis. We now establish a critical role for LAT1 overexpression in enabling the growth of endocrine resistant breast cancer cells.

## Materials and Methods

### Cell lines

LCC1^29^ cells (antiestrogen sensitive) and LCC9^15^ cells (antiestrogen resistant) were obtained from stocks maintained by the Georgetown Tissue Culture Shared Resource. LCC1 and LCC9 cells were cultured in 5% charcoal stripped calf serum (CCS) in phenol red free modified IMEM media (Life Technologies). The MCF7:WS8 cells and T47D cell variants were a gift from Dr. V.C. Jordan at MD Anderson. MCF7:WS8^30^ cells were maintained in 5% fetal bovine serum in modified IMEM media (Life Technologies). T47D:A18 cells and T47D:A18-4HT^16^ cells were grown in 5% fetal bovine serum (FBS) in RPMI (Life Technologies). T47D:C42^17^ cells were grown in 5% charcoal stripped calf serum in phenol red free modified RPMI media (Life Technologies). These cells represent acquired estrogen independence and endocrine therapy resistance in estrogen receptor positive breast cancer. All cells grown in FBS media were estrogen deprived in CCS media for 72 hours before experimental use. All experiments were done in triplicate unless stated otherwise.

### Stable isotope labelling by amino acids in cell culture (SILAC)

MCF7:LCC1 and MCF7:LCC9 cells were double-labelled in presence of heavy (C13) or light (C12) arginine and lysine amino acids. The cells were cultured for at least five doublings before being harvested and snap frozen. Replicates were collected using the label switch approach to assess robustness. MS Bioworks, (Ann Arbor, MI, USA) carried out the SILAC experiments. Label incorporation of more than 98% was confirmed for both the cell lines. The samples were washed with PBS and lysed with RIPA. Ten microgram total protein of light and heavy labelled samples were combined and the combined samples were processed by SDS-PAGE. For each sample, the mobility region was excised into 20 equal sized segments. Each segment was processed by in-gel digestion. Each gel digest was analyzed by nano LC-MS/MS with a Waters NanoAcquity HPLC system interfaced to a ThermoFisher Q Exactive mass spectrometer. Data were processed using MaxQuant version 1.5.3.17 (Max Planck Institute for Biochemistry) that incorporates the Andromeda search engine.

### Pharmacological agents

17β-estradiol (Cat# E-8875) was purchased from Millipore Sigma (MA, USA). 4-hydroxytamoxifen (Cat# 3412) and fulvestrant (Cat# 1047) were purchased from Tocris (Bristol, United Kingdom) and used at pharmacologically relevant concentrations^15,31^. JPH203 (Cat# 406760) was purchased from MedKoo Biosciences Inc. (NC, USA).

### Plasmids and transfections

SLC7A5 siRNA and plasmid DNA were obtained from OriGene. Products used were SLC7A5 (ID 8140) Trilencer-27 Human siRNA and SLC7A5 (NM_003486) Human cDNA ORF Clone. GRP78 (HSP5A) siRNA was purchased from Dharmacon (L-008198-00-0005). Transfections for siRNA used Invitrogen’s Lipofectamine RNAiMAX and for plasmid DNA we used Lipofectamine LTX Plus (ThermoFisher, MA, USA). Cells were treated for 24 hours then refed with fresh growth medium for another 48 hours before collection for protein or growth assay in knockdown experiments. For overexpression experiments, cells were transfected with the appropriate cDNA construct for 4 hours in serum free media before the media was changed to 5% CCS IMEM for an additional 44 hours.

### Crystal violet cell assay

To measure changes in cell growth, 10,000-15,000 cells were plated into each well of a 24-well plate. Treatments were started after 24 hours of seeding (Day 0) and the initial plate was collected as the baseline measurement. At the time of harvest, cells were washed with 1X PBS and rocked in 200 µL crystal violet solution (2.5 g crystal violet, 125 mL methanol, 375 mL water) for 30 minutes. Plates were rinsed in deionized water and allowed to air-dry for 48 hours. Once all time points were collected, citrate buffer was used to extract dye. Analysis of the intensity of staining, which directly reflects cell number, was then assessed at 570 nm using a VMax kinetic microplate reader (Molecular Devices Corp., Menlo Park, CA)^13,32^.

### Western blotting

Total protein was collected in radioimmunoprecipitation buffer (RIPA) with PhosSTOP (Roche Diagnostics, Mannheim, Germany) and Complete Mini protease inhibitor cocktail tablets (EMD Chemicals Inc. San Diego, CA). Quantification was done using Pierce BCA protein assay (Thermo Fischer Scientific) and 20 µg were separated by NuPAGE 4-12% Bis-Tris gel (Invitrogen). Primary antibodies used were LAT1 (Cat #5347S, 1:1,000; Cell Signaling), CD98 (Cat# sc-376815, 1:1,000; Santa Cruz), ER alpha (Cat# sc-543, 1:1,000; Santa Cruz Biotechnology), and β-Actin (Cat# 66009-1-Ig, 1:10,000; Protein Tech). Secondary antibodies used were Anti-rabbit IgG, HRP-linked Antibody #7074 (1:2,000; Cell Signaling) and Anti-mouse IgG, HRP-linked Antibody #7076 (1:2,000; Cell Signaling).

### RNA isolation and qRT-PCR

RNA was isolated using the trizol reagent (Invitrogen, CA, USA) and Qiagen RNeasy mini kit (CA, USA) according to the manufacturer’s instructions. 1 mL of trizol was used per well of a 6-well plate, mixed with 200 uL of chloroform, and incubated at room temperature for 15 minutes. The solution was spun at 15,000 rpm for 15 minutes and the top aqueous layer removed and mixed with an equal volume of 70% ethanol before loading onto the column of the RNeasy kit and processed as described by the manufacturer. Quantification was done using a nano-drop ND-1000 Spectrophotometer. cDNA was made using High Capacity cDNA Reverse Transcription Kit (Thermo Fischer Scientific) to prepare cDNA from 1000 ng RNA. PowerUp SYBR Green Master Mix from Life Technologies was used for qRT-PCR. Primers used were IDT SLC7A5 (FW: CGA GGA GAA GGA AGA GGC G; RV: GTT GAG CAG CGT GAT GTT CC), SLC3A2 (FW: GTC GCT CAG ACT GAC TTG CT; RV: GTT CTC ACC CCG GTA GTT GG), and 36B4 (FW: GTG TTC AAT, GGC, AGC, AT; RV: GAC ACC CTC CAG GAA GCG A). Analysis of the data followed the delta-delta CT method^33^.

### Immunofluorescence staining

10,000-50,000 cells were plated onto glass cover slips 24 hours before treatment. Immunofluorescence experiments were performed on cells after 24 hours of either vehicle or 1 nM 17β-estradiol exposure. Cells were fixed with PBS containing 3.2% paraformaldehyde (Cat# 15714, PA, USA) with 0.2% Triton X-100 (Cat# T8532-500mL, SIGMA, MA, USA) for 5 minutes before being washed with PBS. Cells were incubated in methanol in the −20 ^°^C for 20 minutes. Cells were washed again before being exposed to primary antibody in the presence of an antibody block containing 10% goat serum. Primary antibodies were as described in the western blotting protocol above; the concentration of LAT1 was 1:100 and was 1:50 for CD98. Secondary antibodies used were Alexa Fluor 594 anti-rabbit (Cat# A-11012, Life Technologies) and Alexa Fluor 488, anti-mouse (Cat # A-11001, Life Technologies).

### Cell cycle analysis

Cells were fixed in 75% ethanol and analyzed by FACS analysis (Georgetown Flow Cytometry/Cell Sorting Shared Resource). Cell sorting of GFP-positive cells was done in 5% CCS Media by the Georgetown Flow Cytometry/Cell Sorting Shared Resource then collected for protein or analyzed for cell cycle analysis. Data were acquired using flow cytometry (BD LSRFortessa; BD Biosciences) and data analysis was performed using FCS express 6 software (De Novo Software, Glendale, CA)

### Clinical correlation analyses

We studied only data from invasive ER+ breast cancers that received at least one endocrine therapy (tamoxifen) from the following publicly available datasets (GSE2990^34^, GSE6532-a^35^, GSE6532-p^35^, GSE9195^36^). Data were analyzed as described by Pearce *et al*^37^. For each probe of interest, the dataset was sorted by the normalized expression value in ascending order. Within each sorted sub-dataset, a cursor was set to move up one sample per iteration through the entire dataset starting from the sample with smallest expression value. At each iteration, survival analysis was performed by comparing the samples on either side of the cursor. The resulting statistics including hazard ratio and log rank test p-value provided one measure of significance per possible division in a dataset. In this study, we used 4 independent clinical datasets and 119 genes (probeset_ids). An additional unpublished dataset (GSE46222) was also analyzed and the results are shown in Table 1.

**Table 1:**
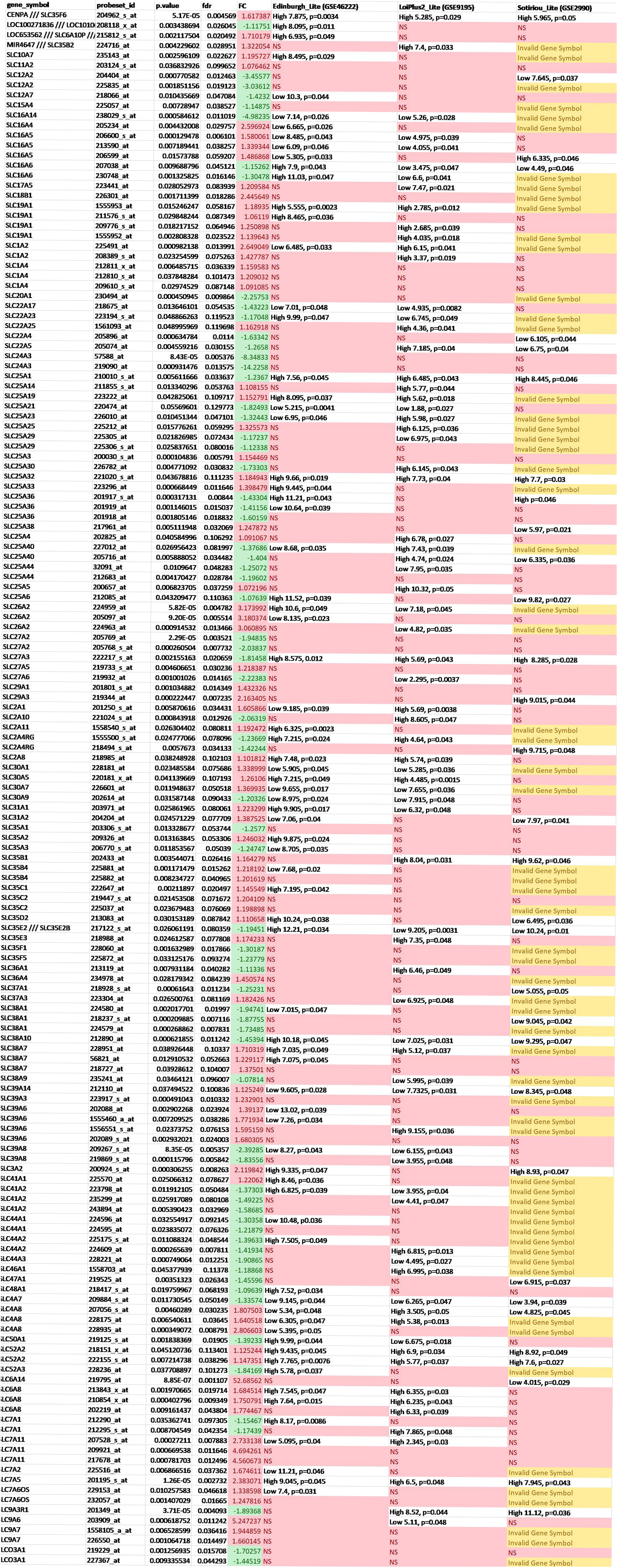
List of significantly differentially regulated genes in LCC9s compared to LCC1s at mRNA analysis and compared with clinical data sets. Three clinical data sets are used: Edinburgh (GSE46222), LoiPlus2 (GSE9195), and Sotiriou (GSE299). For some SLCs multiple probeset_ids exist. For mRNA analysis, p value, FDR, and fold change (FC) exist with FC indicated with positive (red) numbers meaning upregulation and negative (green) numbers meaning downregulation in the LCC9 compared to the LCC1 cells. For the GSE data sets, the direction of high or low expression of the given SLC is indicated for poor prognosis with the p value. Edinburgh and LoiPlus2 utilized the affymetrix HG-U133plus2 chip set while Sotirou used affymetrix HG-U133A chip set resulting in some genes not being included (marked yellow as invalid gene symbol). Non-significant KM data is marked in red as NS.

### Statistical Analysis

ANOVA was used to determine significance (SigmaPlot) with a Dunnett’s *post hoc* test applied when multiple comparisons were made to a common control.

## Results

From microarray analysis of LCC1 and LCC9 mRNA we identified 109 differentially regulated solute carriers (SLCs)^38,39^. Of those 109 solute carriers (SLCs), 55 were associated with poor clinical outcome (Table 1). When mapped onto the proteomic data (TMT and SILAC), three SLCs: SLC2A1, SLC3A2, and SLC7A5 (Table 2) were each upregulated at least 1.5 fold. We had previously studied SLC2A1 (GLUT1) and found that glucose and glutamine uptake are regulated by MYC^13^. Here, we have focused on SLC7A5 (LAT1) and its protein partner SLC3A2 (CD98) to determine their role in endocrine therapy resistance.

**Table 2:**
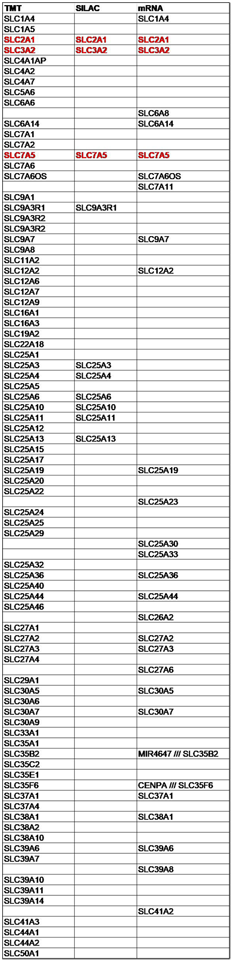
List of significantly upregulated (1.5 fold or more) solute carriers in LCC9 cells compared to LCC1 cells in mRNA, TMT, and SILAC analysis. Red text indicates upregulation in all three data sets.

### LAT1 is regulated by estrogen and upregulated in endocrine therapy sensitive cells

We used MCF7, LCC1, LCC9,^15^ T47D:A18, T47D:C42, and T47D:4HT^17^ cells as models to study the role of LAT1 in endocrine resistance. From the three analyses, we found that LAT1 was significantly upregulated in the endocrine resistant LCC9 compared with endocrine sensitive LCC1 cells. Previously, LAT1 (SLC7A5) mRNA has been reported to be induced by estrogen in MCF7 cells^40^. We confirmed this observation for LAT1 protein in MCF7 cells (Figure 1A) and found a similar induction in LCC1 and T47D:A18 cells (Figure 1A&B). Endocrine resistant LCC9, T47D:4HT, and T47D:C42 cells expressed significantly higher basal levels of total LAT1 protein than their respective (sensitive) parental cell lines. Since these data infer estrogen regulation of LAT1 by the estrogen receptor (ER; ESR1), we used ENCODE to analyze ChIA-PET data from MCF7 cells^41,42^. Using the integrative genomics viewer, we found an ER-occupied site on the LAT1 gene (Figure 1C; Supplemental Figure 1 for whole gene)^43^.

**Figure 1:**
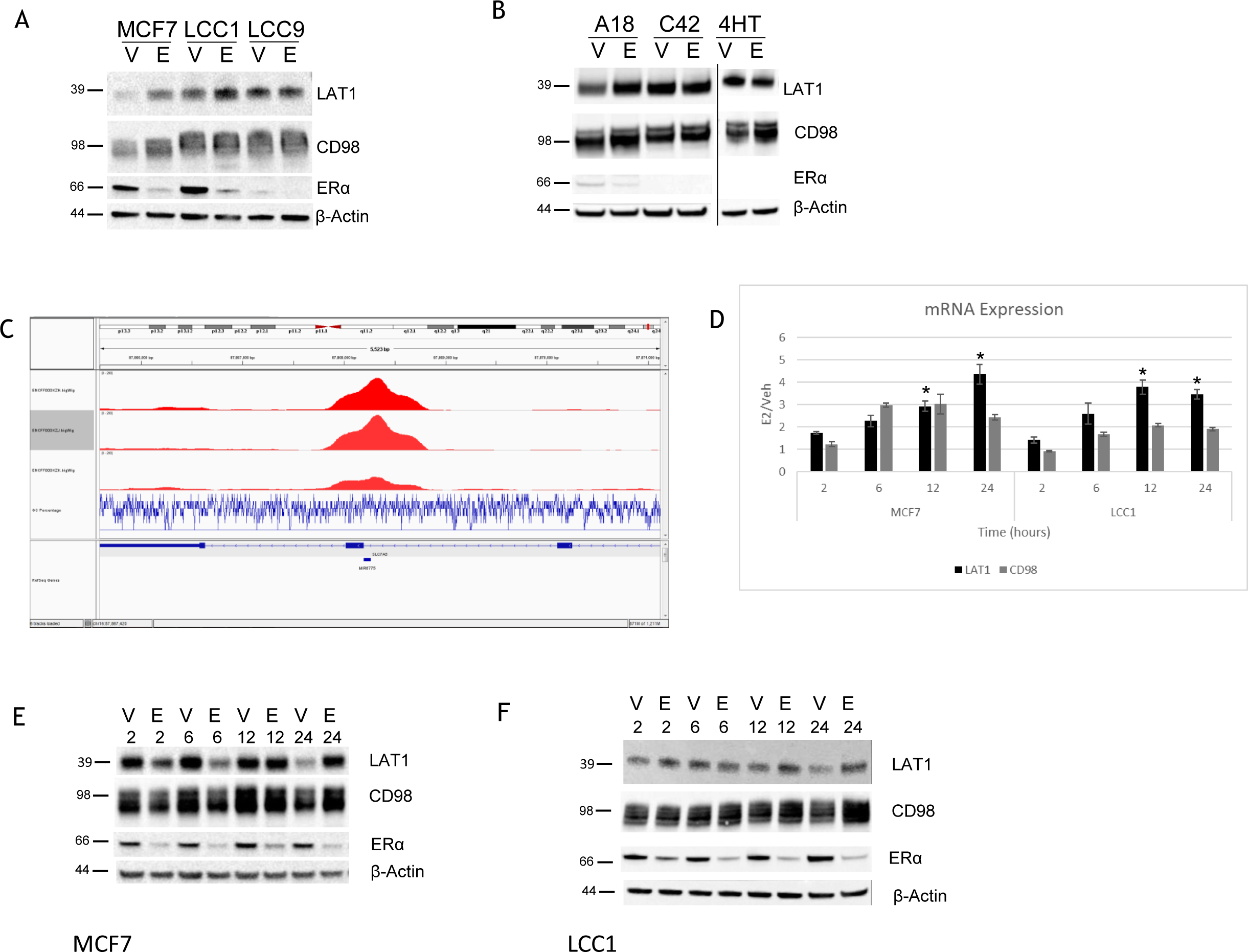
LAT1 is estrogen regulated in endocrine therapy sensitive cell lines. A) LAT1 and CD98 are upregulated in endocrine therapy resistant cells (LCC9s) compared to sensitive cell lines (MCF7, LCC1s). B) This upregulation was observed in T47D:C42 and T47D:4HTs compared to parental T47D:A18s. C) ESR1 binding on the LAT1 gene. D) Increasing time of estrogen (1nM) increases LAT1 and CD98 mRNA in both MCF7 and LCC1 cells. E) MCF7 and F) LCC1 cells show increased protein levels of LAT1 with estrogen treatment.

Endocrine sensitive cells doubled their LAT1 protein expression when treated with estrogen for 24 hours. The LAT1 mRNA and protein expression increased in response to estrogen over time in both MCF7 and LCC1 cells (Figure 1D-F). After 12 to 24 hours of estrogen treatment, LAT1 mRNA levels were significantly upregulated (p<0.05). In contrast to the endocrine sensitive models, LCC9 and T47D:C42 cells both showed an increase in basal LAT1 protein expression but no further increase of LAT1 was seen after 24 hours of estrogen treatment.

### E2 regulation of LAT1 is lost in endocrine resistant LCC9 cells

Since we observed an estrogen-induced increase of LAT1 protein and mRNA expression in sensitive cell lines, we next studied an extended estrogen time course treatment in resistant cells. Levels of LAT1 mRNA and protein did not change in response to estrogen with increased time of treatment in the resistant LCC9 cells (Figure 2A-B). This loss of regulation did not affect either basal expression levels or membrane subcellular co-location of the LAT1 and CD98 proteins. Protein co-localization was measured in immunofluorescence experiments where both MCF7 and LCC9 cells were treated with either vehicle or estrogen (Figure 2C-D respectively).

**Figure 2:**
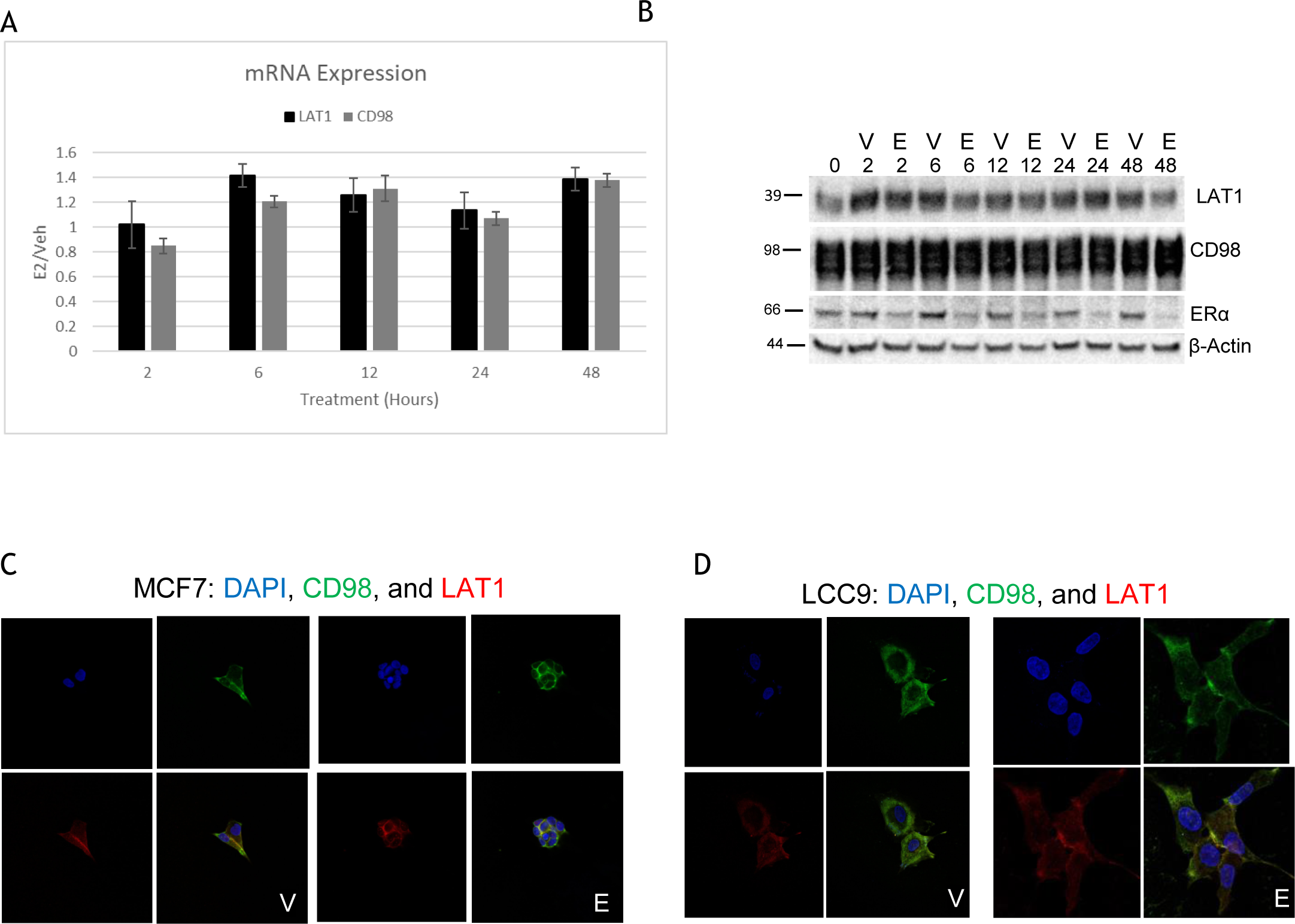
LAT1 is not upregulated by estrogen in endocrine therapy resistant LCC9s. A) LAT1 nor CD98 are significantly changed at the mRNA level with estrogen treatment. B) Western blot analysis also showed no difference at the protein level. C) MCF7 and D) LCC9 cells look similar in immunofluorescent images as LAT1 and CD98 co-localize in both cell lines.

### LAT1 is differentially expressed in response to endocrine therapy treatment

To determine the effect of endocrine therapies on LAT1 expression, combinations of estrogen and either tamoxifen or fulvestrant were used to determine how LAT1 was regulated in response to estrogen treatment. MCF7 (Figure 3A), LCC1 (Figure 3B), and LCC9 cells (Figure 3C) express both mRNA and protein for LAT1 and CD98. LAT1 expression was significantly increased in response to estrogen or tamoxifen in the sensitive models but unchanged in resistant models. Fulvestrant decreased LAT1 expression in MCF7 and LCC1 cells treated with E2 or tamoxifen, suggesting that ER inhibition negatively affects LAT1 expression. Estrogen alone, tamoxifen alone, and estrogen and tamoxifen cotreatment each significantly increased LAT1 mRNA (p<0.05) in both sensitive models. In the MCF7 cells, addition of fulvestrant with tamoxifen did not return LAT1 mRNA levels fully to baseline (upregulation p<0.05). In the MCF7 models the classical estrogen-regulated GREB1 mRNA was not upregulated in response to tamoxifen but increased in response to estrogen (Supplemental Figure 2A-B). GREB1 mRNA was unchanged in the LCC9 cells in response to endocrine treatments (Supplemental Figure 2C). In the T47D models, the same trend was seen between the endocrine sensitive T47D:A18s and resistant T47D:C42 and T47D:4HT cells (Supplemental Figure 3).

**Figure 3:**
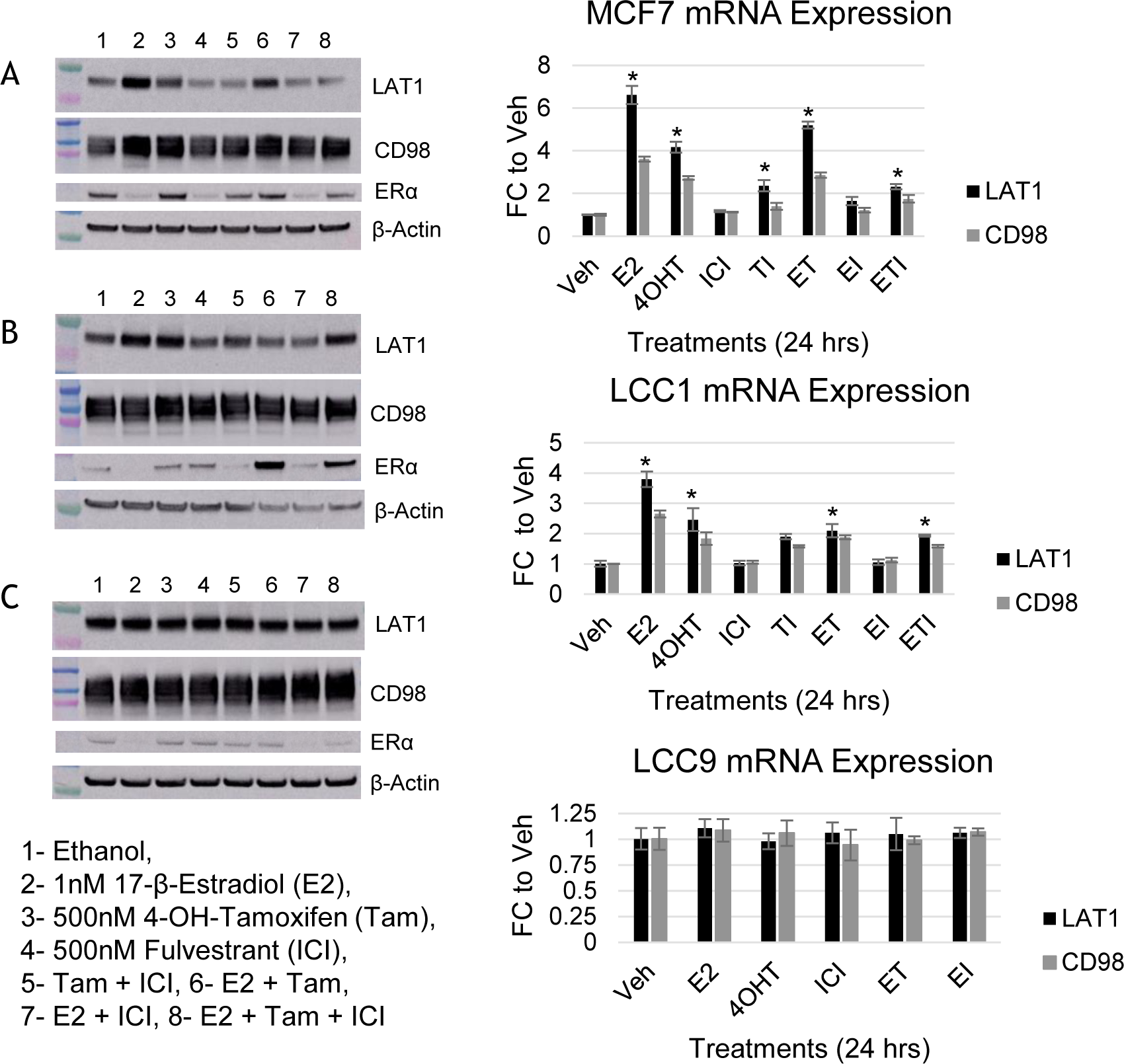
Endocrine therapies differentially change LAT1 expression in sensitive but not resistant cells. A) MCF7, B) LCC1, and C) LCC9 cell lines show differential LAT1 and CD98 protein or mRNA expression with endocrine therapy treatment for 24 hours. Endocrine therapy sensitive cells upregulate LAT1 and CD98 in response to estrogen and tamoxifen treatment.

### LAT1 inhibition restricts cell growth and induces G1 arrest

Since LAT1 had increased basal expression and lost estrogenic regulation in resistant cells, we targeted LAT1 function using JPH203, a tyrosine analog and selective inhibitor of LAT1 function^44,45^. We applied a time-(3 to 6 days) and dose-dependent study design (12.5 - 50 μM) to determine how MCF7 and LCC9 cells respond to JPH203 treatment (Figure 4A-B). Growth was significantly inhibited by ~50% with 50 μM JPH203 in both cell lines in the presence or absence of estrogen (p>0.05). The effect of JPH203 treatment increased when we reduced the concentration of essential amino acids in the media in a dose-dependent manner (Supplemental Figure 4). We also used two individual siRNAs to knock-down LAT1 expression. 72 hours after transfection, LAT1 protein expression was decreased by 40-60% as confirmed by Western blot hybridization (Figure 4C-D). Cell growth was significantly decreased with two individual siRNAs targeting LAT1 (Figure 4E, p<0.05) after 3 or 6 days compared with control. To determine how LAT1 inhibition affected cell cycle distribution, we performed cell cycle analysis of MCF7 and LCC9 cells treated with either 50 μM JPH203 or with siLAT1. While JPH203 inhibition did not change cell cycle phase distribution of the MCF7 or LCC9 cells, treatment with siLAT1 decreased the proportion of cells in S phase (Figures 4F-H). Puromycin is an inhibitor of global protein synthesis and can be used to assess translation by treating cells with a high dose followed by a western blot^46^. LCC9 cells treated with siLAT1 followed by puromycin treatment exhibited a decrease in global protein translation (Figure 4I). Targeting LAT1 either pharmacologically or genetically was effective in reducing growth of the resistant cells.

**Figure 4:**
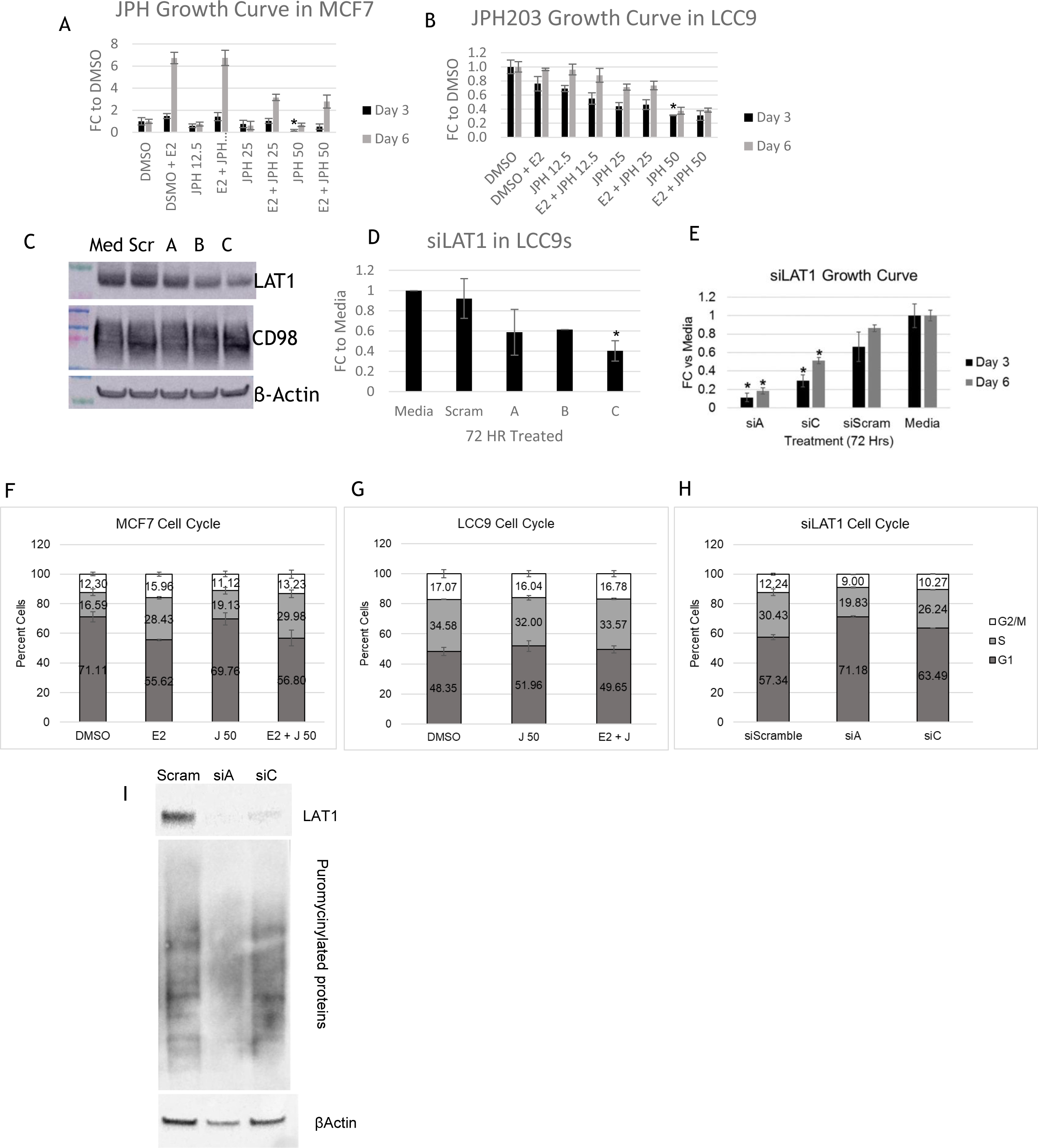
LAT1 inhibition restricts MCF7 and LCC9 proliferation. A) MCF7 and B) LCC9 growth curves when treated with increasing doses of JPH203 for 3 and 6 days. C) siRNA targeting of LAT1 which is quantified in D) was more effective. E) Growth curve of siLAT1 cells showed a decrease in in cell growth consistent for 3 or 6 days. Cell cycle analysis of F) MCF7 and G) LCC9 with JPH203 did not yield significant results, however H) siRNA knockdown of LAT1 in LCC9s reduced S phase. I) Western blot of puromycinylated proteins showed a reduction in global protein translation with LAT1 knock down after 72 hrs.

### Overexpression of LAT1 increases S phase and global protein translation

MCF7 and LCC1 cells were transfected with plasmids containing either a GFP-empty vector or GFP-LAT1 cDNA. Overexpression of LAT1 protein was confirmed in MCF7 and LCC1 cells by measuring protein expression (Figure 5A and 5B respectively). Fluorescence imaging of the GFP tag (Figure 5C) also confirmed plasmid expression. Puromycin treatment of transfected cells showed an increase in global protein translation (Figure 5D). In addition to increased global translation in the sensitive cells, we observed an increased trend for cells to be in S phase in both MCF7 and LCC1 cells (Figure 5E).

**Figure 5:**
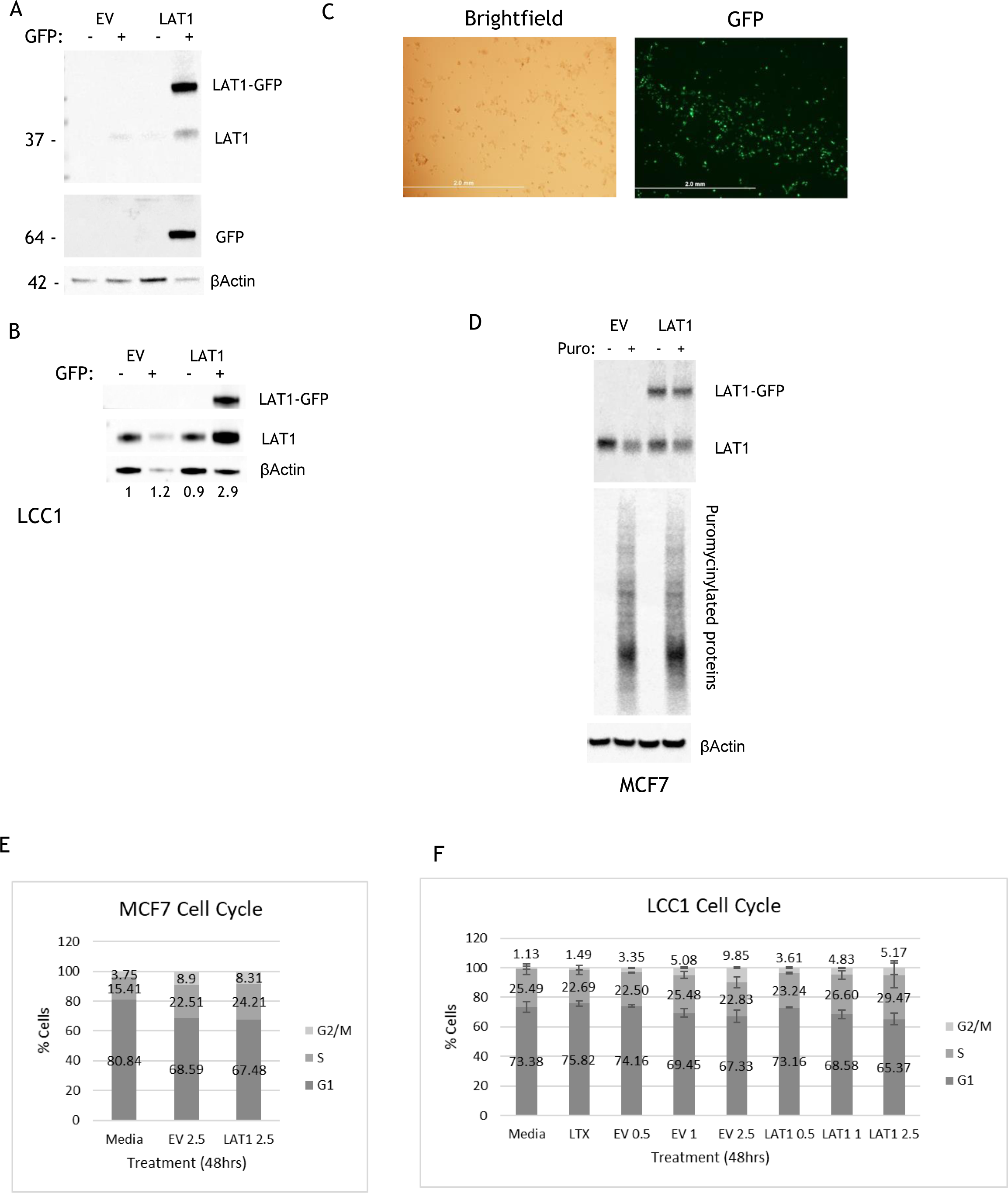
LAT1 Overexpression leads to proliferative advantage in MCF7 and LCC1s. LAT1 plasmid was transfected into cells with a GFP tag. A) MCF7 cells and B) LCC1 cells were sorted for GFP positivity confirmed LAT1 overexpression through western blot. C) microscopy image shows LAT1 overexpression in MCF7s. D) MCF7 cells were treated with puromycin to show increased global protein translation with LAT1 overexpression. Cell cycle analysis shows a trend increase of S phase in both E) MCF7 and F) LCC1 cells.

### Autophagy increases with LAT1 inhibition

Increased autophagy is a feature of endocrine resistant cells^12,47^ that may cooperate with increased nutrient scavenging by SLCs to support the restoration of metabolic homeostasis. Autophagic flux can be estimated by measuring the expression of two key proteins: LC3 and p62^48^. Apoptosis can be evaluated by western blot hybridization of the cleavage of poly(ADP-ribosyl) polymerase (PARP)^49^. Expression of both the LC3 and p62 proteins was increased following siRNA knockdown of LAT1 (Figure 5A). LC3 expression increased but p62 did not decrease. These data are consistent with an induction of autophagy but incomplete autophagic flux. Neither PARP cleavage nor phosphorylation of eIF2a was observed, suggesting that apoptosis and the PERK pathway within the unfolded protein response (UPR) are not required for this process. Knockdown of GRP78 in LCC9 cells (BiP; controls all three pathways within the UPR including PERK) produced a non-significant increase in LAT1 protein expression, whereas LAT1 mRNA expression was significantly increased (Figure 6C and D). These data imply either an increased rate of GRP78 protein turnover or a delay in increasing mRNA translation; determining the precise mechanism is beyond the scope of the current study.

**Figure 6:**
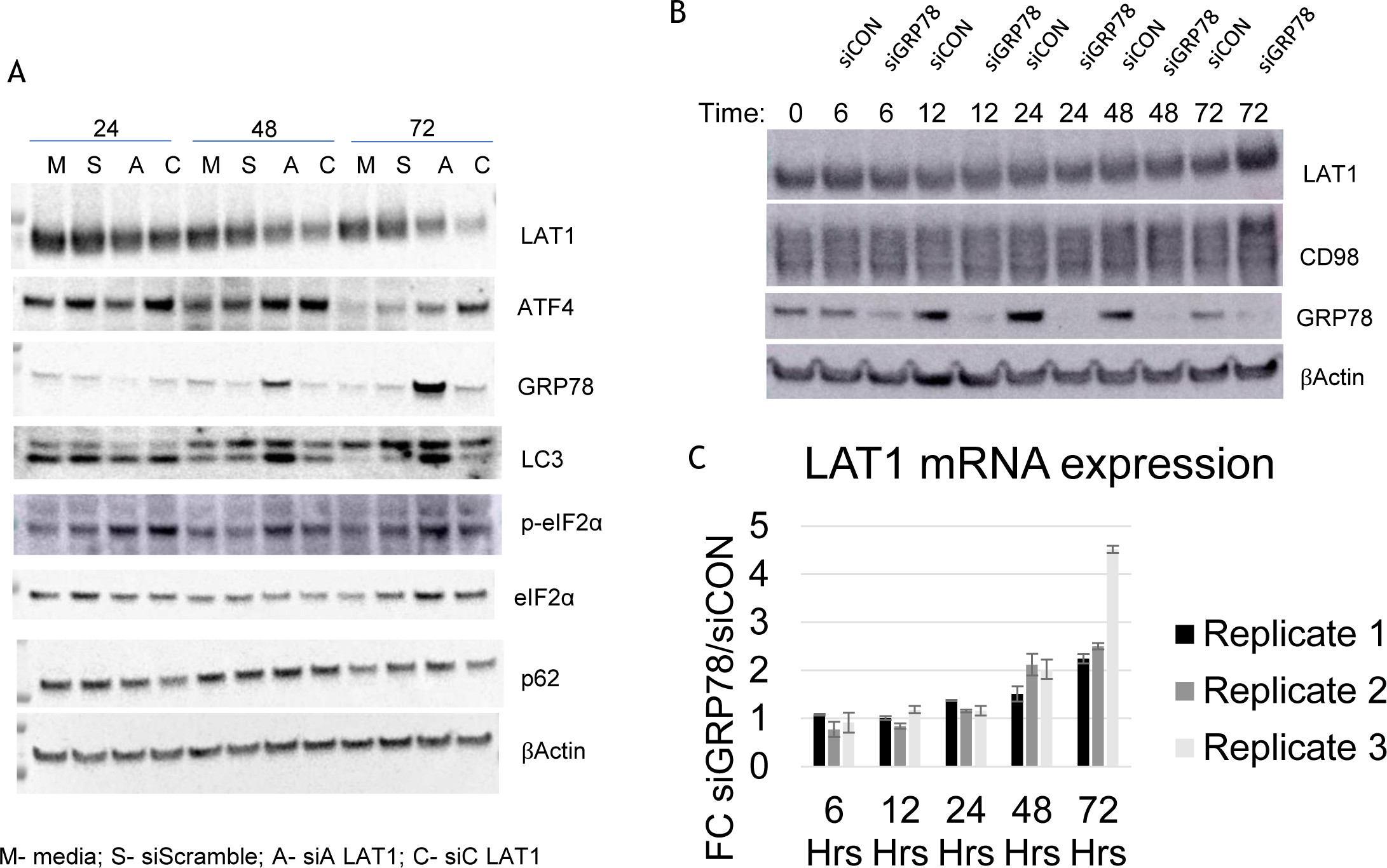
Autophagy initiates but does not complete with LAT1 inhibition. A) markers for autophagy and the unfolded protein response show autophagic flux. B) knockdown of GRP78 shows an increase of LAT1 protein after 72 hours. C) The mRNA levels of LAT1 increase with GRP78 knockdown (one replicate is stronger than the other).

### Higher LAT1 expression correlates with poor clinical outcome

To determine the clinical relevance of LAT1 in endocrine-treated ER+ breast cancer, we established the association of LAT1 mRNA expression with clinical outcomes in four gene expression data sets (Figure 7, see Materials and Methods). We studied only invasive ER+ breast cancers that received at least one endocrine therapy. Data sets were analyzed as described by Pearce et al.^37^ Higher LAT1 expression correlates with a poor disease-free survival (From KM plots GSE2290 p=0.007, GSE6532-a p=0.005, GSE6532-p p=0.037, GSE9195 p=0.01 and Table 2).

**Figure 7:**
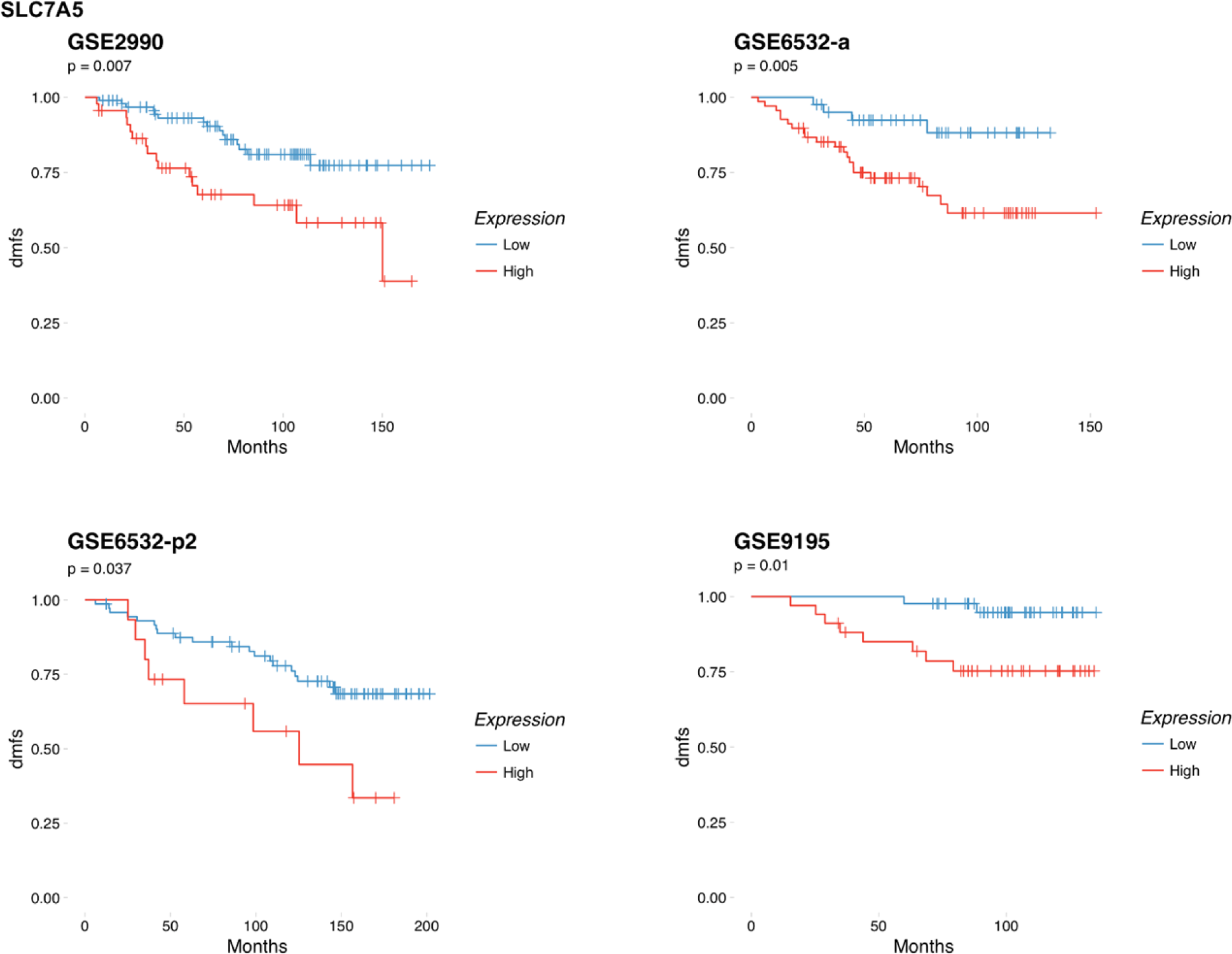
Clinical data sets confirm increased LAT1 expression correlates with poor disease-free survival.

## Discussion

While tamoxifen and fulvestrant are effective endocrine therapies^3^, further research is needed to prevent or overcome the development of resistance. Endocrine therapy resistance, particularly in advanced disease, is a major clinical challenge for patients and their physicians. Matched sensitive and resistant cell lines are useful tools to study changes in cell processes as endocrine therapy resistance develops. By performing mRNA, TMT, and SILAC analyses of differentially regulated genes, we identified several key players associated with the development of acquired resistance (Table1). SLC7A5 (LAT1) was significantly upregulated in the LCC9 compared with the LCC1 cells in all three analysis. This observation led to our focus on LAT1 to determine its role in the development or maintenance of endocrine therapy resistance.

LAT1 has been proposed as a biomarker for progression in breast cancers^50^. However, the role of LAT1 in the context of endocrine therapy responsiveness is unknown. Our study shows that LAT1 overexpression in endocrine resistant breast cancer cells contributes to their survival and growth. For example, we establish that LAT1 mRNA and protein expression are increased in endocrine resistant breast cancer cells compared with their genetically related but endocrine sensitive counterparts. LAT1 is reported to be estrogen regulated^40^; we confirmed this observation using ER positive MCF7 and T47D breast cancer cells. Constitutive activation of the ER is one component of endocrine resistance that results in the dysregulation of a number of downstream genes^51^. Notably, in endocrine resistant cells the basal expression of LAT1 was higher and its estrogenic regulation was lost. A drug-induced reduction of amino acid uptake in sensitive cells could lead to metabolic stress and ultimately cell death. Resistant cells must find a way to address this limitation. Upregulation of SLCs such as LAT1 could improve a cell’s ability to scavenge nutrients from the tumor microenvironment, a function that is critical for cell survival^52^. LAT1 is responsible for the uptake of leucine and tyrosine for protein synthesis or as intermediates to enter the TCA cycle^19,53,54^. Increased LAT1 expression has been reported in several cancers including prostate cancer^22^, pleural mesothelioma^24^, multiple myeloma^25^ and non-small cell lung cancer^26^. Since homozygous knockout of LAT1 in embryonic lethal^28^, LAT1 is critical for growth and survival.

While LAT1 is under estrogen regulation in MCF7 and LCC1 cells, this is lost in the resistant cells. LAT1 expression was increased by tamoxifen treatment in both MCF7 and LCC1 cells; this increase was reduced by fulvestrant. These observations are likely reflective of the partial agonist activity of tamoxifen and further imply that LAT1 expression is under estrogenic regulation. MYC is also under estrogenic regulation and we have shown that MYC can regulate glucose and glutamine through the unfolded protein response in endocrine resistant cells^13^. LAT1 upregulation in endocrine resistance may cooperate with MYC-induced increases in glucose and glutamine metabolism to contribute to cell survival in the face of the stress induced by endocrine therapies.

Targeting solute carriers has not been widely explored in breast cancer. JPH203 was less effective than a targeted siRNA knockdown to restrict cell growth and induce G1 arrest. However, free tyrosine, leucine, and phenylalanine in media likely influenced the efficacy of JPH203; these and other free amino acids also may be accessible within the tumor microenvironment. siRNA knock-down of LAT1 also initiated autophagy and decreased global protein translation in endocrine resistant LCC9 cells. The latter could be controlled by activation of the UPR^56^, which can also regulate autophagy^12,47^. Inhibiting LAT1 would reduce amino acid uptake that could activate autophagy in an attempt to restore metabolic homeostasis. However, LAT1 inhibition lead to an initiation of autophagy but flux did not complete and cell death occurred. Knocking down GRP78, the primary regulator of the unfolded protein response (UPR)^57^, increased LAT1 expression after 72 hours. It is likely that the uptake of amino acids and the ability of UPR to regulate global protein translation are connected, perhaps by activating features of the UPR.

JPH203 showed antineoplastic activity and safety for biliary tract and colorectal cancer in the Phase I clinical trial reported by Okana et. al.^58^ While LAT1 inhibitors have not yet been tested in breast cancer patients, the drug appears to be well tolerated. Targeting LAT1 limits the amount of amino acids, particularly leucine and tyrosine, that can enter the TCA cycle or maintain the production of new proteins^27,40^. Using JPH203 in combination with endocrine therapies and/or mTOR inhibitors^59^ could prove beneficial. For example, the combination of JPH203 and mTOR inhibitors could result in decreased amino acid uptake and protein translation to restrict tumor cell growth. Further exploration into the metabolic fate of the increased uptake of pre-formed amino acids could provide useful insights into the metabolic adaptations required to maintain endocrine resistance. Imaging of leucine or tyrosine with positron emission tomography (PET)^60^ could be clinically informative as a potential biomarker of endocrine responsiveness in ER+ breast tumors.

Finally, we show that LAT1 overexpression is consistently associated with poor relapse free survival in four independent clinical datasets from ER+ patients that received an endocrine therapy. This observation is consistent with other reports that LAT1 may be an indicator of poor prognosis^25,61^. Taken together with the mechanistic outcomes reported here, further study of LAT1 and its role in endocrine therapy resistance may lead to novel therapeutic alternatives to improve overall survival for patients.

## Acknowledgements

This work was supported by Public Health Service Awards U54-CA149147, U01-CA184902 (R Clarke) and Department of Defense Breast Program W81XWH-18-1-0722 (R Clarke) and Lombardi Comprehensive Cancer Center Support Grant (CCSG) NIH P30 CA051008. We thank Karen Creswell and Dan Xun for their help at the Flow Cytometry Shared Resource at Georgetown-Lombardi Comprehensive Cancer Center. The views and opinions of the author(s) do not reflect those of the US Army or the Department of Defense.

---

**Supplemental Figure 1:**
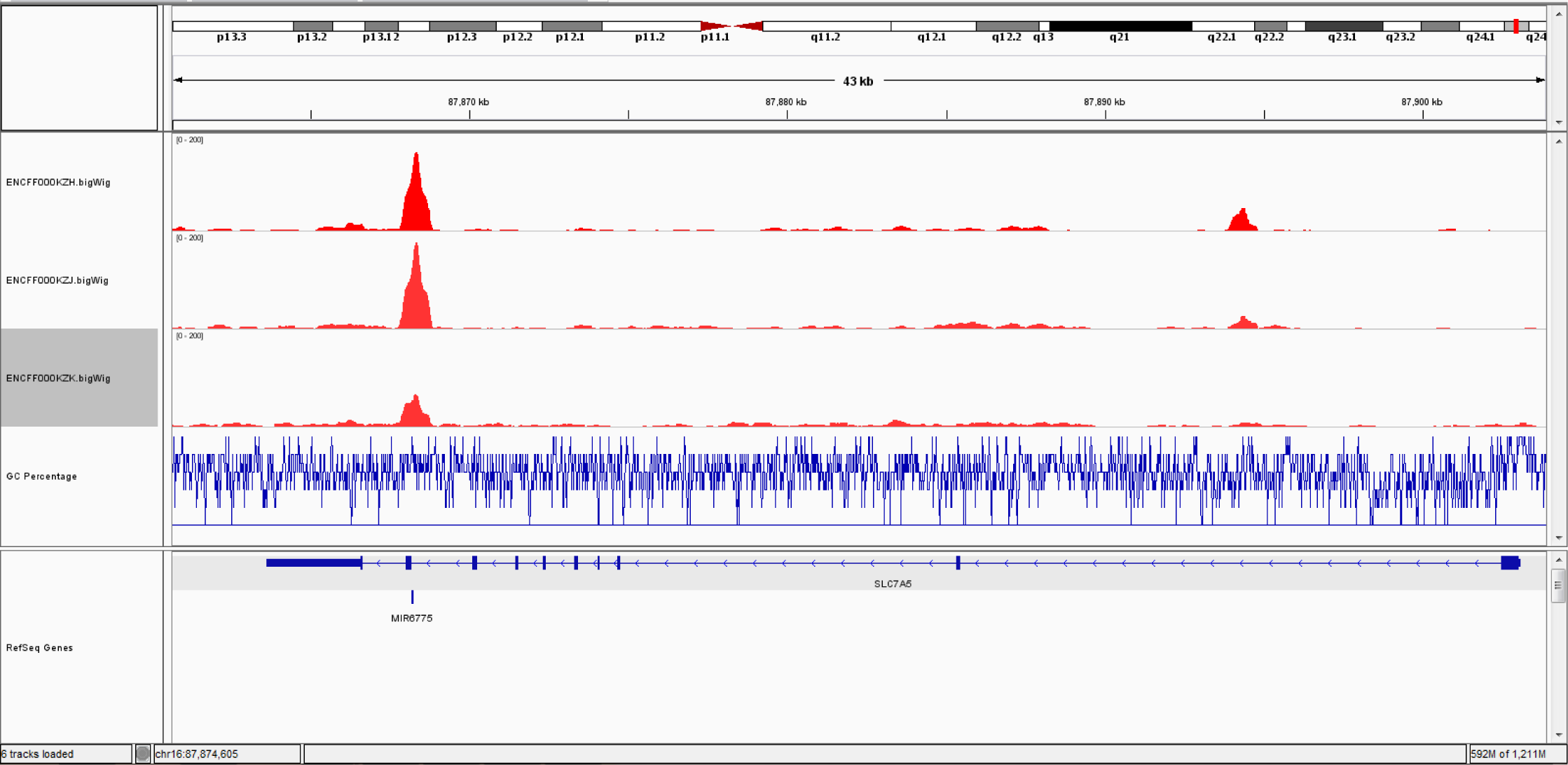
ESR1 protein binds to an early portion of the LAT1 gene in MCF7 cells as shown by ChIA-PET.

**Supplemental Figure 2:**
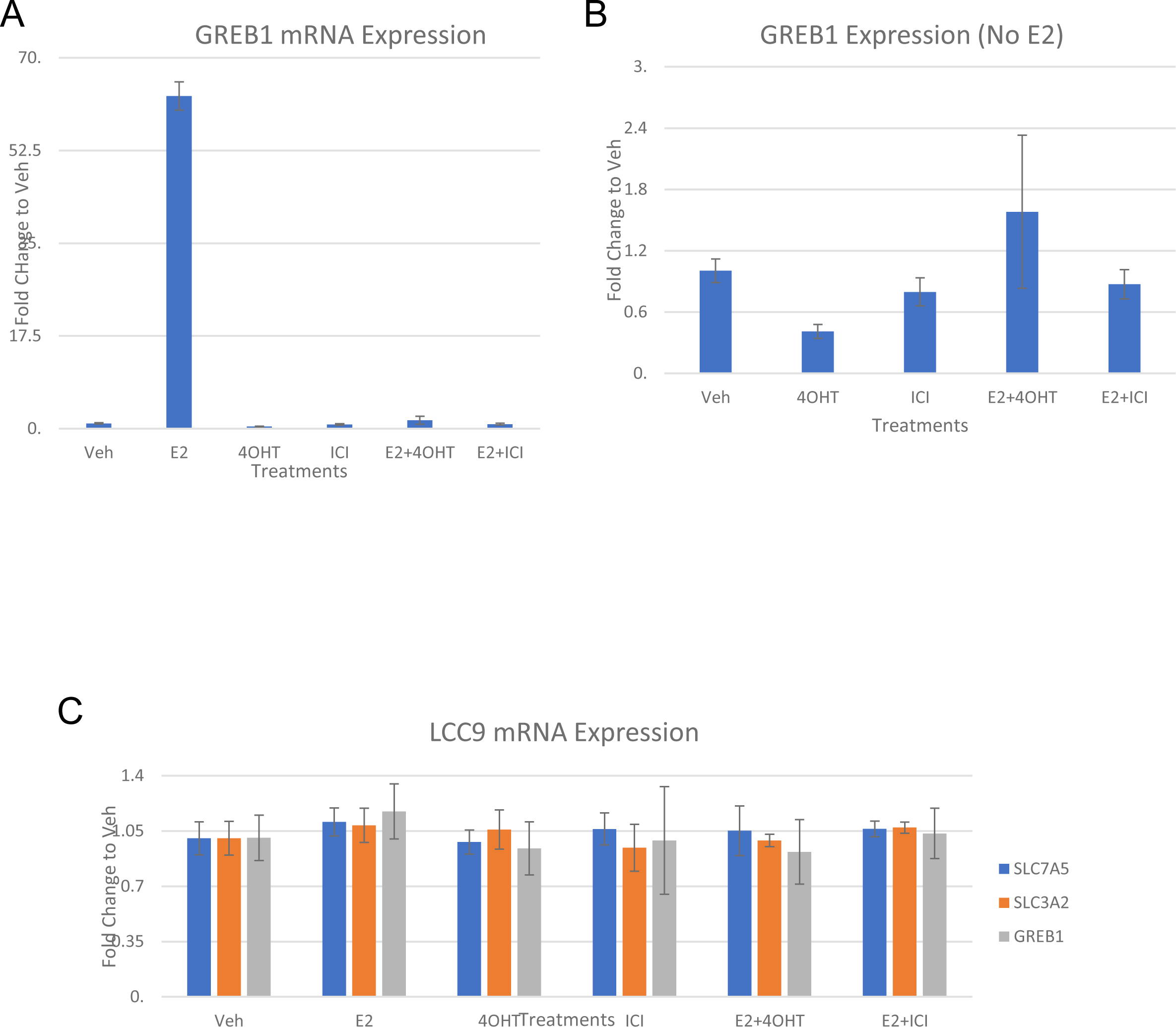
GREB1 is classically expressed in response to endocrine therapy treatment in MCF7 but not LCC9s. A) GREB1 mRNA expression increases with estrogen treatment. B) GREB1 mRNA does not change in response to endocrine therapy treatment showing classical estrogen regulation which is lost in C) LCC9 cells.

**Supplemental Figure 3:**
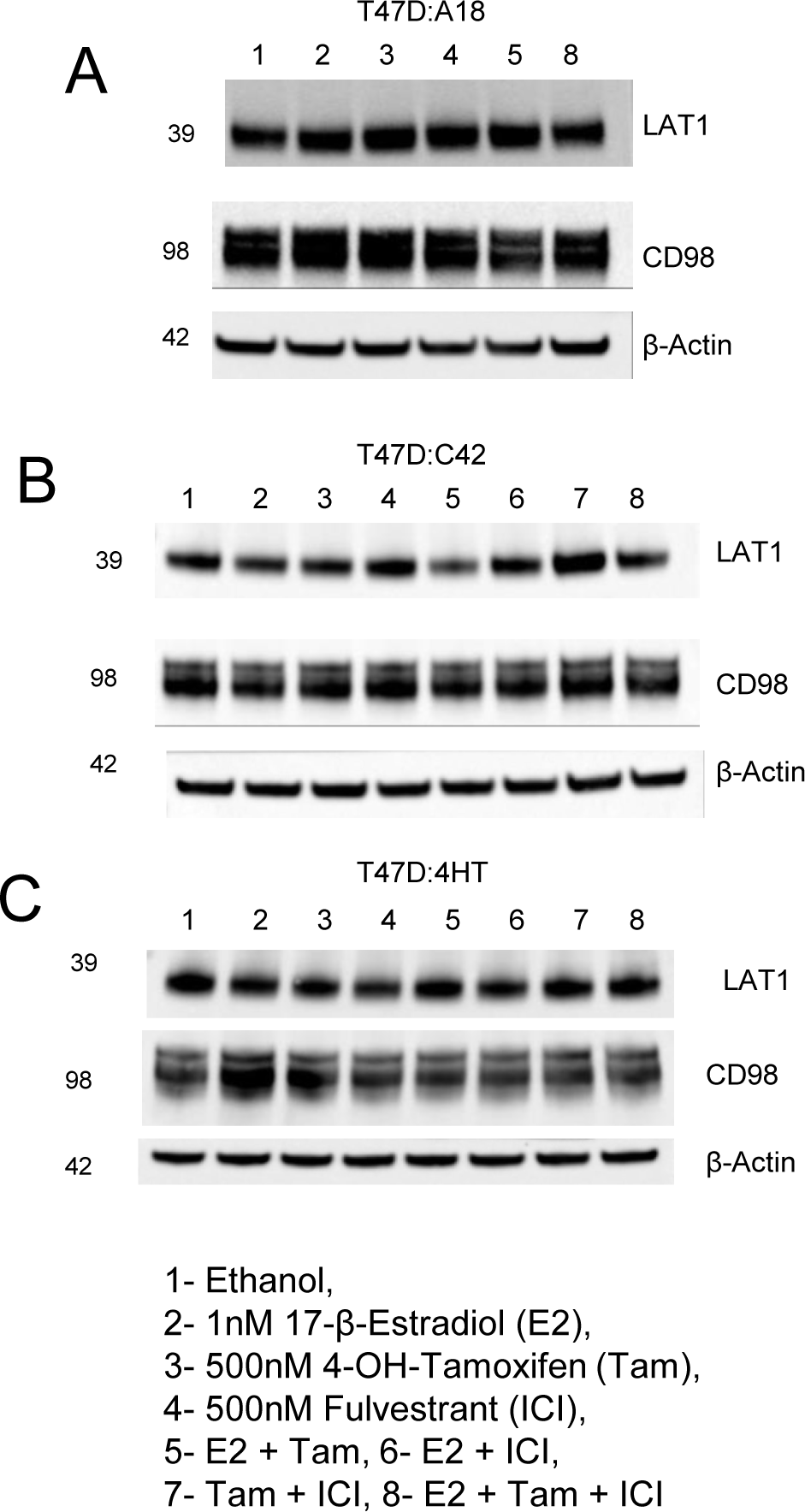
Endocrine therapy treatment modulates LAT1 expression in A) endocrine therapy sensitive T47D:A18s but not endocrine therapy resistant B) T47D:C42s nor C) T47D:4HTs.

**Supplemental Figure 4:**
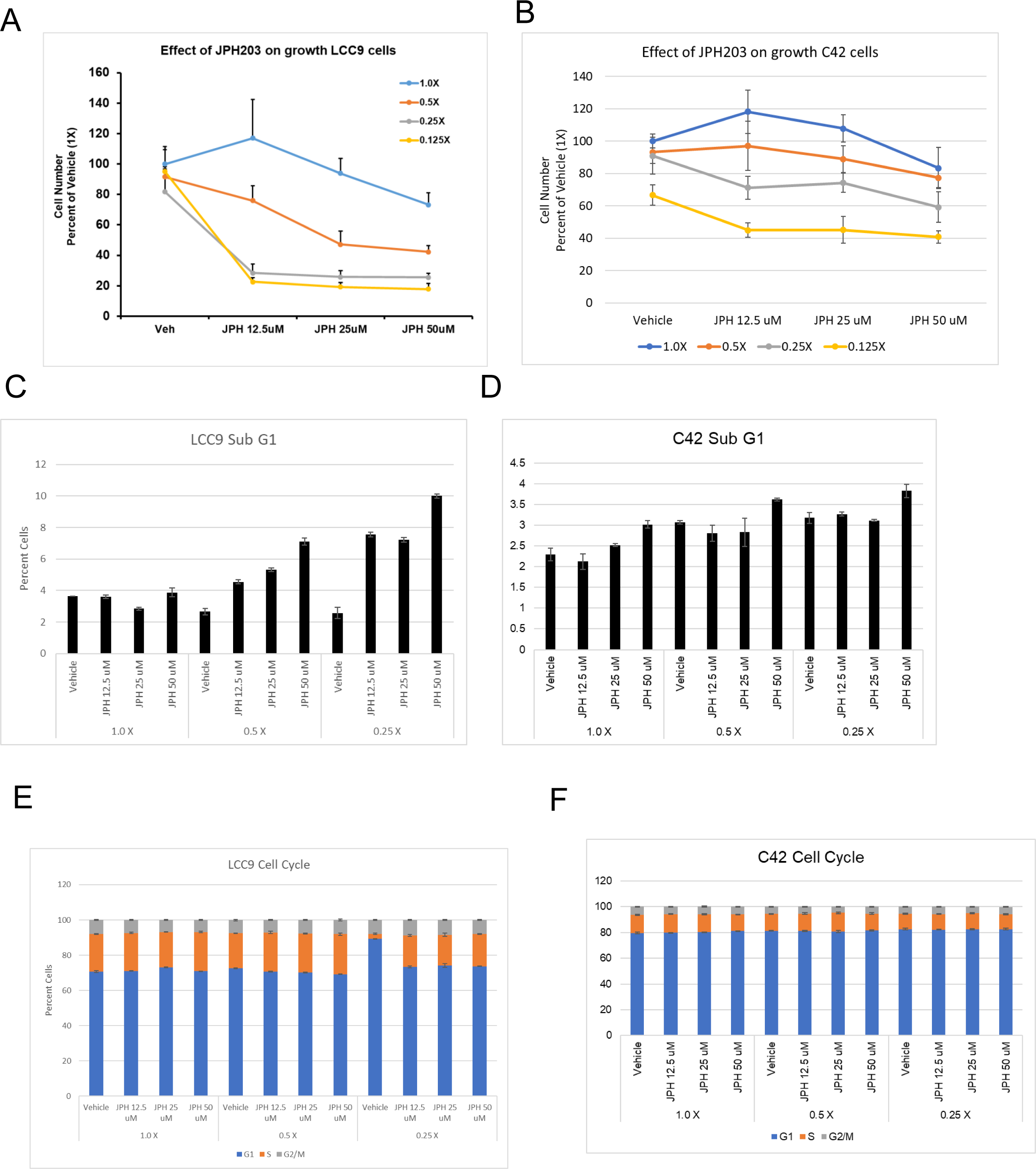
Depleting essential amino acids in the media enhances growth arrest by JPH203 in both LCC9s and C42s. A) LCC9s or B) T47D:C42s were cultured in essential amino acid deplete media and treated with indicated concentrations of JPH203 resulting in increased efficacy. C) LCC9 and D) T47D:C42 cells had an increase in Sub G1 when analyzed by flow cytometry and E-F) both showed a decrease in S phase.

